# Host metabolic reprogramming of *Pseudomonas aeruginosa* by phage-based quorum sensing modulation

**DOI:** 10.1101/577908

**Authors:** H. Hendrix, M. Kogadeeva, M. Zimmermann, U. Sauer, J. De Smet, L. Muchez, M. Lissens, I. Staes, M. Voet, J. Wagemans, P-J. Ceyssens, J-P. Noben, A. Aertsen, R. Lavigne

## Abstract

The *Pseudomonas* quinolone signal (PQS) is a multifunctional quorum sensing molecule of key importance to the *P. aeruginosa* metabolism. We here describe that the lytic *Pseudomonas* bacterial virus LUZ19 targets this population-density-dependent signaling system by expressing <u>q</u>uorum <u>s</u>ensing-associated acyl<u>t</u>ransferase (Qst) during early infection. Qst interacts with a key biosynthesis pathway enzyme PqsD, resulting in decreased metabolites levels of PQS and its precursor 2-heptyl-4(1H)-quinolone. The lack of a functional PqsD enzyme impairs the normal LUZ19 infection but is restored by external supplementation of 2-heptyl-4(1H)-quinolone, showing that LUZ19 exploits PQS to successfully achieve its infection. A functional interaction network, which includes enzymes of the central carbon metabolism (CoaC/ThiD) and a novel non-ribosomal peptide synthetase pathway (PA1217), suggests a broader functional context for Qst, which blocks *P. aeruginosa* cell division. Qst represents an exquisite example of intricate reprogramming of the bacterium, which may be exploited towards antibiotic target discovery for this bacterial pathogen.

## Introduction

Bacteriophages are the most abundant biological entities on Earth, affecting bacteria and even global ecosystems through microbial mortality (Suttle, 2007), horizontal gene transfer (McDaniel *et al.*, 2010) and metabolic reprogramming (De Smet *et al.*, 2016). Due to intimate co-evolution with bacteria, phages efficiently and extensively adapt the host physiology. This hijacking primarily takes place immediately after infection and is observed at all levels of the cellular metabolism (De Smet *et al.*, 2017). At the molecular level, this hijacking is achieved through i.a. protein-protein interactions of mainly early phage proteins that inhibit, activate or functionally alter specific host proteins (Hauser *et al.*, 2012; Roucourt and Lavigne, 2009). These early proteins can also include metabolic enzymes, of which the genes are termed <u>a</u>uxiliary <u>m</u>etabolic <u>g</u>enes (AMGs) (Thompson *et al.*, 2011). AMGs generally encode for proteins which complement rate-limiting enzymes to compensate for imbalances arising during infection, especially in nucleotide biosynthesis (Enav *et al.*, 2014; Miller *et al.*, 2003) and energy production (Sharon *et al.*, 2011; Sullivan *et al.*, 2006). However, emerging evidence supports a more general model of phage-directed host reprogramming in which AMGs influence nearly the entire central carbon metabolism (Hurwitz *et al.*, 2013; Thompson *et al.*, 2011).

This host takeover is not self-evident and is preceded by a struggle for cellular control between the phage and the host (Samson *et al.*, 2013). A recently identified strategy is the use of quorum sensing systems by *Pseudomonas aeruginosa* to activate its defense mechanisms against phage predation. Overall, the risk of phage infection is the highest at high cell density. Known phage resistance mechanisms regulated by quorum sensing are the reduction of phage receptors on the cell surface (Høyland-Kroghsbo *et al.*, 2013) and the activation of the prokaryotic adaptive CRISPR-Cas immune system (encoded by the **C**lustered **R**egularly **I**nterspaced **S**hort **P**alindromic **R**epeats loci and **C**RISPR-**as**sociated (Cas) genes) (Patterson *et al.*, 2016; Høyland-Kroghsbo *et al.*, 2017). More generally, quorum sensing influences the cell’s physiological state as to better protect it against phage attack (Qin *et al.*, 2017).

In *Pseudomonas aeruginosa*, four main quorum sensing systems have been identified to date: the N-<u>a</u>cyl<u>h</u>omoserine lactones (AHL)-dependent systems LasI/LasR and RhlI/RhlR (Pesci *et al.*, 1997), the <u>i</u>ntegrated <u>Q</u>uorum sensing <u>s</u>ignal (IQS) system AmbBCDE/IqsR (Lee *et al.*, 2013) and the *<u>P</u>seudomonas* <u>q</u>uinolone <u>s</u>ignal (PQS) system PqsABCDH/PqsR (Dubern and Diggle, 2008). These signaling systems are believed to regulate the expression of up to 10% of *Pseudomonas* genes, including self-regulation (autoinduction), cross-regulation between the different quorum sensing systems and regulation of bacterial virulence and biofilm formation (Schuster *et al.*, 2003; Williams and Camara, 2009). Remarkably, about half of the quorum sensing-regulated genes still have an unknown function (Schuster *et al.*, 2003). Besides cell density, the quorum sensing systems can also be influenced by environmental factors such as phosphate-limiting conditions (Lee *et al.*, 2013) and oxidative stress (Häussler and Becker, 2008). For example, the PQS system plays a key role in protecting a micro-colony by both stimulating the destruction of damaged sister cells and rescuing undamaged members from oxidative stress by lowering their metabolic activity (D’Argenio *et al.*, 2002; Häussler and Becker, 2008).

Phages have evolved mechanisms to interfere with their host’s quorum sensing systems, possibly (i) to overcome the CRISPR-Cas mediated phage resistance, (ii) to oppose quorum sensing-mediated downregulation of specific phage receptors (Høyland-Kroghsbo *et al.*, 2013), (iii) to control biofilm formation (Pei and Lamas-Samanamud, 2014), (iv) to influence the quorum sensing-regulated cell physiological state which naturally protects bacteria against phage attacks (Qin *et al.*, 2017), (v) to alter bacterial behavior via response regulators to enhance phage production, or (vi) to utilize the quorum sensing molecule as a carbon source for phage progeny (Müller *et al.*, 2014). In the *Clostridium difficile* phage phiCDHM1, three AMGs from the host accessory gene regulator (agr) quorum sensing system (AgrB, AgrC and AgrD) were proposed, while response regulators associated with this system have been predicted in three *Pseudomonas* phages (Hargreaves *et al.*, 2014). Furthermore, *Iodobacte*r phage fPLPE encodes a putative acylhydrolase, for which its bacterial homolog degrades the AHL signal molecules (Leblanc *et al.*, 2009), and *Vibrio* phage VP882 encodes a quorum sensing receptor homolog that guides the lysis-lysogeny decision (Maxwell, 2019; Silpe and Bassler, 2019). So far, no mechanisms in phages are known to interfere with PQS signaling. Nevertheless, a recent observation by our group hints at an important role of the PQS signal in the host response to phage infection (Blasdel *et al.*, 2018; De Smet *et al.*, 2016).

Phage LUZ19 is a podovirus of the *Autographivirinae* subfamily, for which the genome organization has been intensely studied (Lavigne *et al.*, 2013). We here show that a protein expressed during early infection, Gp4 (termed Qst, ‘<u>q</u>uorum <u>s</u>ensing-associated acyl<u>t</u>ransferase), modulates the PQS levels in the host, resulting in toxicity and metabolic reprogramming. This function is associated with acyltransferase activity and is achieved by a complex interaction network, in which Qst interacts with enzymes of both central carbon metabolism and two signaling molecule biosynthesis pathways: i) the well-known quorum sensing molecule PQS and ii) a predicted non-ribosomal peptide (the PA1221 cluster; Gulick, 2017). The interacting protein of the latter pathway is also able to neutralize the antibacterial effect of Qst, hinting a phage-mediated effect on cell signaling. The further exploration of these novel phage-host interactions may lead to insights that could aid in the development of new antibacterial therapies targeting quorum sensing (Fetzner, 2015).

## Results

### An early-expressed phage LUZ19 protein shows antibacterial activity in *P. aeruginosa*, which is complemented by the hypothetical protein PA1217

During a screen for growth-inhibitory ORFans in *P. aeruginosa* PAO1-infecting phages, the early expressed protein Gp4 of phage LUZ19 (referred to hereafter as Qst) was identified (Wagemans *et al.*, 2014). Upon induction of Qst, *P. aeruginosa* growth was completely abolished (Figure 1A) and a filamentous growth type was observed (Figure 1B). Interestingly, this toxicity is rather specific, as *E coli* MG1655 cells grew normally after episomal expression of Qst. Qst has a predicted molecular weight of 13 kDa, has no conserved domains and is only found in *Kmvvirus* phages.

**Figure 1.**
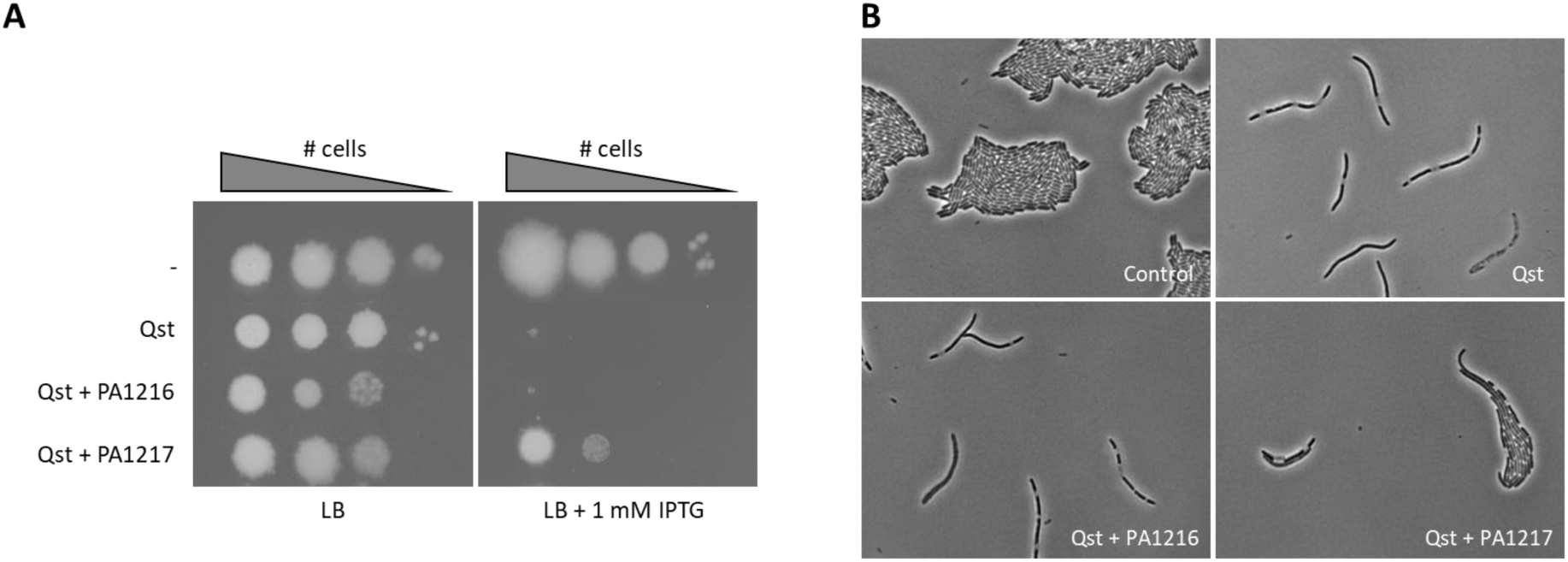
Antibacterial effect of the early phage protein Qst on *P. aeruginosa* PAO1 growth and its partial complementation by the hypothetical protein PA1217. (A) Hundredfold serial dilutions of *P. aeruginosa* PAO1 strains harboring the *qst* gene (second row), both *qst* and *PA1216* genes (third row), and both *qst* and *PA1217* genes (bottom row) were spotted on medium with (right) or without (left) IPTG induction, together with a negative control (empty vector construct, top row). (B) *P. aeruginosa* PAO1 morphology after Qst expression (top right), and both Qst and PA1216/PA1217 expression (bottom), together with a control (empty vector construct, top left). Microscopic recording of *P. aeruginosa* cells after growth for 5 h in the presence of 1 mM IPTG.

To study the mechanisms underlying this toxicity, we performed a complementation screen to identify bacterial proteins that can alleviate this toxic effect upon overexpression. The twenty confirmed positive clones containing random *P. aeruginosa* PAO1 genome fragments all matched to a single locus in the *Pseudomonas* genome, with five different fragments that overlapped by 3,227 bp (position 1,316,424-1,319,651 bp, minus-strand) encoding two putative ORFs, PA1216 and PA1217. After individual cloning into a *Pseudomonas* expression vector and expression of PA1216 or PA1217 in the Qst expressing *P. aeruginosa* PAO1 strain, a spot test and microscopic analysis revealed that overexpression of PA1217 can partially complement the toxic effect of the phage protein on the host cells (Figure 1A, B).

### Qst binds to the predicted CoA acetyltransferase region of PA1217

To test whether the antibacterial effect of Qst was due to direct protein-protein interaction with PA1217, we performed both *in vitro* and *in vivo* interaction analyses. First, a native gel mobility shift assay with an equal amount of recombinant PA1217 and an increasing amount of Qst protein displayed a small but distinct shift in PA1217 migration in presence of Qst. Excising and loading the shifted band from the native gel on a denaturing gel, proved the presence of both PA1217 and Qst in the shifted band (Figure 2A). Furthermore, an ELISA using PA1217 as bait and Qst as prey further validated this interaction, as an interaction was observed independently from the quantity of Qst and the ratio between Qst and PA1217 (Figure 2B). Together, these results confirmed that Qst directly interacts with PA1217 *in vitro*.

**Figure 2.**
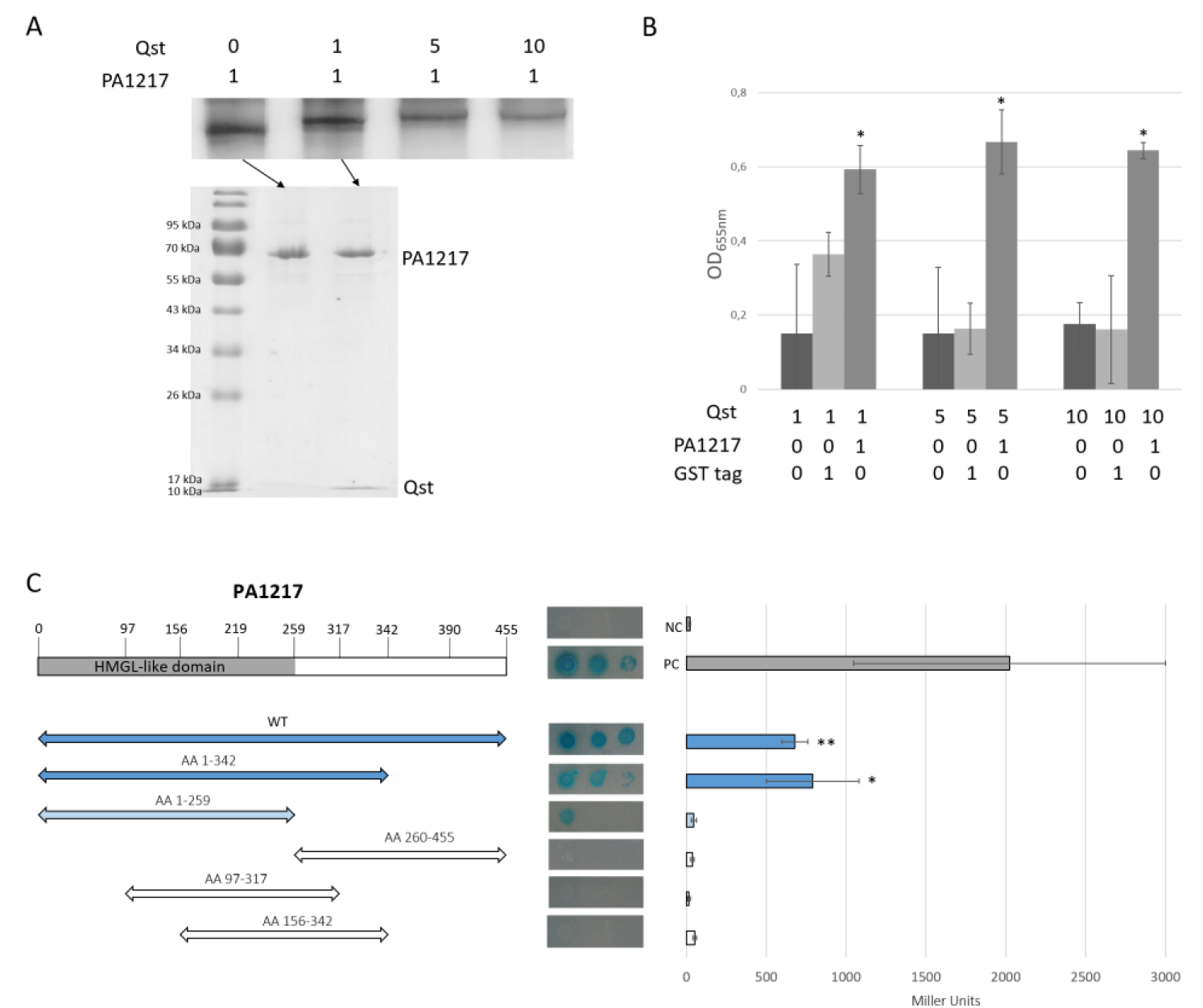
Interaction analyses of Qst and the *P. aeruginosa* hypothetical protein PA1217. (A) Native gel mobility shift assay using an equal amount of PA1217 and an increasing amount of Qst (top). The band in lane 1 and the shifted band in lane 2 were excised and analyzed via SDS-PAGE (bottom). The numbers indicate the relative amount of Qst and PA1217 (4 pmol). (B) Enzyme-linked immunosorbent assay using an equal amount of GST-tagged PA1217 (10 pmol) as bait and an increasing amount of His-tagged Qst as prey. As negative controls, no bait protein and GST tag were used. The numbers indicate the relative amount of Qst, PA1217 and GST tag. The Y-axis indicates the absorbance measured at OD_655nm_ after 10 min, using Anti-Mouse IgG antibody conjugated to HRP and 1-Step Slow TMB-ELISA substrate. Error bars represent standard deviation and *P*-values (compared to no bait protein and GST tag separately) were calculated using the Student’s t-test (n=3), *p<0.05. (C) Bacterial two-hybrid assay in which the T18 domain is N-terminally fused to Qst and the T25 domain is N-terminally fused to PA1217 or a fragment of PA1217. Fragments of the PA1217 protein are shown on the left. Blue and white arrows indicate fragments with a positive and no signal, respectively. Interactions were visualized by a drop test on minimal medium (pictures in the middle), and quantified by measuring the β-galactosidase activity, which are indicated in Miller Units (graph at the right; Figure 2-figure supplement 1). Non-fused T25 and T18 domains were used as a negative control, and the leucine zipper of GCN4 was used as a positive control. Error bars represent standard deviation and *P*-values (compared to both combinations with empty counterparts) were calculated using the Student’s t-test (n=3), *p<0.05, **p<0. 01.

Follow-up *in vivo* validation of this interaction and specification of the interaction site using bacterial two-hybrid showed significant (p<0.05) positive reactions for Qst with both the full PA1217 protein and its N-terminal part (residues 1-342), containing a catalytic HMGL (hydroxymethylglutaryl-coenzyme A lyase)-like domain. However, the HMGL-like domain alone (residues 1-259) did not appear to bind Qst, perhaps due to lack of proper folding of this fragment (Figure 2C, Figure 2-figure supplement 1).

### Qst and its interaction partner PA1217 both show acyltransferase activity

The *P. aeruginosa* protein PA1217 is predicted as a probable 2-isopropylmalate synthase, containing an HMGL-like domain (residues 1-259; Pfam, E-value: 1.2e-61). Isopropylmalate synthases belong to the DRE-TIM metallolyase superfamily, which share a TIM-barrel fold, an HXH divalent metal binding motif and a conserved active site helix (Casey *et al.*, 2014). In *P. aeruginosa* one functional homolog is known, LeuA, which catalyzes the first step in leucine biosynthesis. To verify the 2-isopropylmalate synthase activity, acetyl-Coenzyme A to Coenzyme A (CoA) conversion was assayed *in vitro* (Roucourt *et al.*, 2009). Although both the positive control LeuA and PA1217 showed transferase activity, the activity of PA1217 was markedly lower, suggesting the absence of its natural substrate (Figure 3). ^*1*^*H-NMR* analysis confirmed this assumption, as products were only formed by LeuA, while PA1217 merely consumed acetyl-CoA (Figure 3-figure supplement 1). PA1217 is embedded in an uncharacterized non-ribosomal peptide synthetase (NRPS) cluster (Gulick, 2017), controlled by quorum sensing regulation and producing a potential novel cell-to-cell signaling molecule, which may explain the lack of 2-isopropylmalate synthase activity as it possibly functions in the modification of the non-ribosomal peptide and has therefore alternative substrate specificity. Also in other NRPS clusters, homologs of 2-isopropylmalate synthases are identified (Jenul *et al.*, 2018; Rouhiainen *et al.*, 2010).

**Figure 3.**
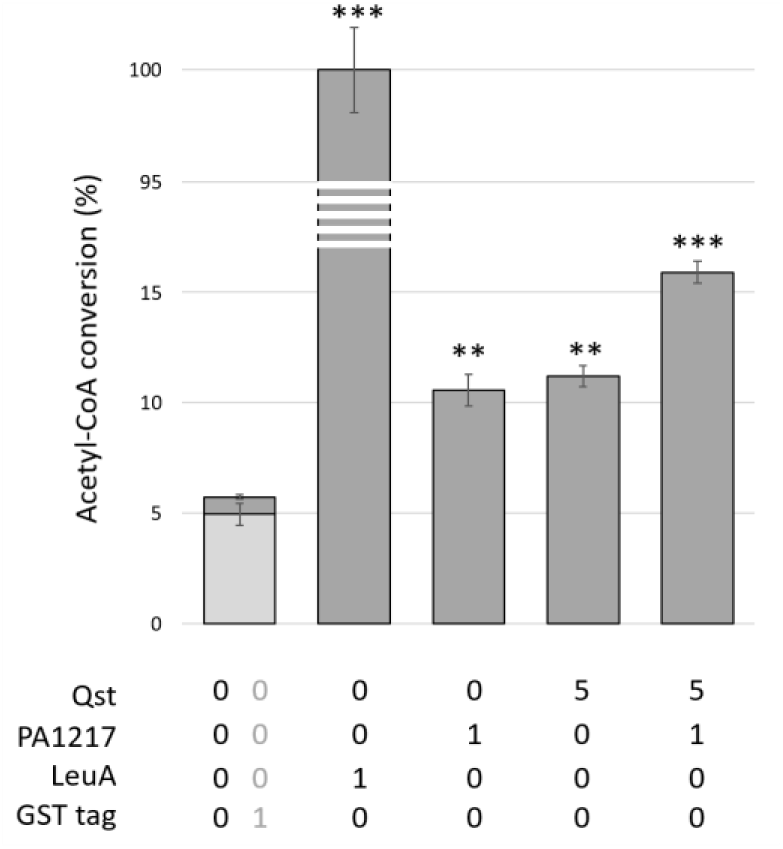
CoA acetyltransferase activity assay. The acetyl-CoA transferase activity was measured using the Ellman’s reagent (DTNB) after 30 min incubation at 37°C, and expressed relative to the activity of the *P. aeruginosa* 2-isopropylmalate synthase LeuA. Error bars represent standard deviation and *P*-values (compared to both controls) were calculated using the Student’s t-test (n=3), **p<0.01, *** p<0.001. The numbers below indicate the relative amount of the proteins, i.e. 1 µg of PA1217 and 5 µg of Qst were used in the reaction mixtures.

Based on the interaction analysis, we tested a possible inhibitory effect by Qst on the enzymatic activity of PA1217. Surprisingly, however, we observed that Qst was also able to consume acetyl-CoA, and a Qst/PA1217 combination showed additive enzymatic activity (Figure 3, Figure 3-figure supplement 1). These data suggest that the small early-expressed phage protein Qst interferes with acetyl-CoA metabolism in the cell.

### Untargeted metabolomics reveals alterations in the central carbon metabolism and the *Pseudomonas* quinolone signal (PQS) pathway

To investigate the function of Qst during phage infection, we employed untargeted metabolomics to identify host metabolic pathways which are either directly or indirectly affected by Qst. We compared metabolite levels between *P. aeruginosa* PAO1 expressing recombinant Qst and the wild-type strain, the strain overexpressing PA1217 or the one expressing both Qst and PA1217. In total, the level of 1,140 known metabolites (from the KEGG database (Kanehisa and Goto, 2000)) were monitored before and at various time points after Qst expression (15, 30, 45, 60, 90 and 120 min).

Upon Qst expression, only nine metabolites changed significantly (abs(log2(FC)) > log2(1)) and p-value < 0.05) in at least a single time point of the experiment compared to the wild type and showed a significant change over time (Page’s trend test: false discovery rate (FDR) < 0.05), as shown on the heatmap of fold-changes (Figure 4A).

**Figure 4.**
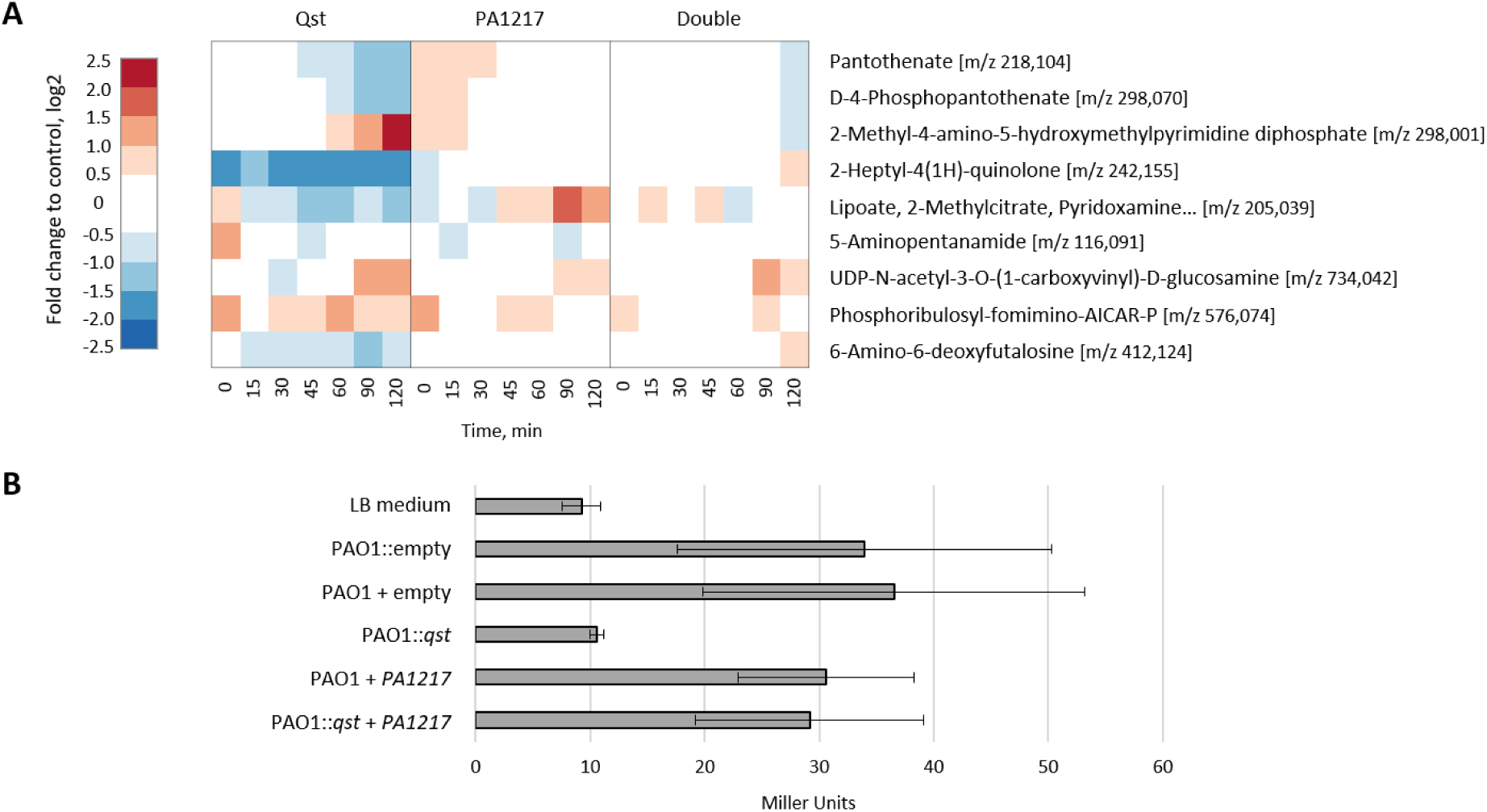
Impact of Qst on metabolite levels in *P. aeruginosa*. (A) Heatmap of significantly changed metabolites after recombinant Qst expression. Only metabolites that meet the following criteria are shown: Page FDR < 0.05, and at least one time point with an abs(log2(FC)) > log2(1)) and p-value < 0.05 (both compared to the wild type). Levels of selected metabolites in a *P. aeruginosa* PAO1 strain overexpressing PA1217, and in a strain both expressing Qst and PA1217 (double) are shown as well. Fold changes represent the means of three biological replicates. (B) PQS bioassay. Cell lysates of 1 mM IPTG-induced, stationary growing *P. aeruginosa* PAO1 strains containing an empty pUC18mini construct (PAO1::empty), an empty pME6032 vector (PAO1 + empty), a pME6032 vector with *PA1217* gene (PAO1 + *PA1217*), a pUC18mini construct with *qst* gene (PAO1::*qst*), and a pUC18mini construct with *qst* gene and a pME6032 vector with *PA1217* gene (PAO1::*qst* + *PA1217*) were added to *P. aeruginosa* PAO1-R1 (pTS400) cultures. The experiment was quantified by measuring the β-galactosidase activity, which are indicated in Miller Units. Error bars represent standard deviations (n=3).

These significantly changed metabolites include pantothenate and 4-phosphopantothenate, which are precursors for the biosynthesis of both cofactors CoA and 4-phosphopantetheine. Lipoate, another metabolite that is significantly reduced upon Qst expression, functions like CoA and 4-phosphopantetheine as an acyl carrier cofactor (Hazra *et al.*, 2009; Spalding and Prigge, 2010), showing that Qst indeed affects acyltransferase metabolism. This reduction in cofactor precursors could be the result of potential overconsumption of acyl carrier factors (indirect effect) or a potential overactivation of biosynthesis enzymes (through direct interaction). Within this context it is interesting to note that 4-phosphopantetheine is also an essential cofactor for peptidyl carrier proteins, like the NRPS PA1221 belonging to the same secondary metabolite biosynthesis gene cluster as PA1217 (Mitchell *et al.*, 2012). Furthermore, the CoA biosynthesis pathway is known to influence the thiamine biosynthesis pathway, which could explain the increase of the cofactor thiamine pyrophosphate precursor 2-methyl-4-amino-5-hydroxymethylpyrimidine diphosphate (Enos-Berlage and Downs, 1997).

Another interesting observation is the clear decrease in quorum sensing molecule 2-heptyl-4(1H)-quinolone (HHQ) upon Qst expression (Figure 4A). This molecule is synthesized by the gene products of *pqsABCD* and is a precursor of the quorum sensing molecule 2-heptyl-3-hydroxy-quinolone (PQS), which is also decreased (log2(FC) > log2(1) and p-value < 0.07) in all time points after Qst expression (Figure 4-Source Data 1). However, it should be noted that both changes are already present prior to Qst induction, which might be the result of background transcription levels (Choi *et al.*, 2005). Nevertheless, an additional decrease in metabolite levels of HHQ is observed 45 min after induction (Figure 4-figure supplement 1). Furthermore, the metabolite level pattern of PQS in the double expression mutant suggests an initial decrease by Qst, which is subsequently complemented by PA1217 (Figure 4-figure supplement 1). Also, the most abundant quinolone produced from the PQS biosynthesis pathway, 2,4-dihydroxyquinoline (DHQ) (Zhang *et al.*, 2008), is less abundant in *P. aeruginosa* expressing Qst compared to the strains overexpressing PA1217 and expressing both Qst/PA1217 (Figure 4-figure supplement 1). DHQ can bind to the PQS specific transcription regulator PqsR (similar to HHQ and PQS) and is therefore thought to also play a role in *P. aeruginosa* pathogenicity (Gruber *et al.*, 2016). To confirm the reduced levels of PQS upon Qst expression, a PQS bioassay with the *P. aeruginosa* PAO1-R1 (pTS400) *lacZ* reporter strain (Van Houdt *et al.*, 2004) revealed a reduced reaction compared to the other strains. This indeed indicates a lower concentration of PQS present in the cell lysate (Figure 4B).

### Qst interacts with both quorum sensing-associated secondary metabolite and central metabolism pathways

Based on the observed influence of Qst on a range of metabolic pathways, we hypothesized that Qst physically interacts with additional enzymes beyond PA1217. To identify such additional interaction partners, we performed an *in vitro* pull-down using immobilized His-tagged Qst protein as bait and crude *Pseudomonas* cell lysate as prey. The elution fractions showed additional protein bands compared to the control sample (Figure 5A). We analyzed both samples with ESI-MS/MS and identified a total of 104 proteins, including Qst (total spectral count = 29). Interestingly, three enzymes which were not present in the control sample but were pulled-down when the cell lysate was incubated with Qst, could be directly associated to the observed metabolic changes: CoaC, ThiD, and PqsD (Figure 5-Source Data 1). Upon recombinant Qst expression in *P. aeruginosa* the levels of 4-phosphopantothenate, the substrate of CoaC (phosphopantothenoylcysteine synthase/4’-phospho-N-pantothenoylcysteine decarboxylase) decreased, whereas the levels of 2-Methyl-4-amino-5-hydroxymethylpyrimidine diphosphate, the product of ThiD (phosphomethylpyrimidine kinase) increased, suggesting an overactivation of these enzymes. The third pulled-down enzyme, PqsD (3-Oxoacyl-(acyl carrier protein) synthase III) is a key enzyme of the PQS biosynthesis pathway, which can explain the observed decrease in PQS, HHQ and DHQ levels (Figure 4A).

**Figure 5.**
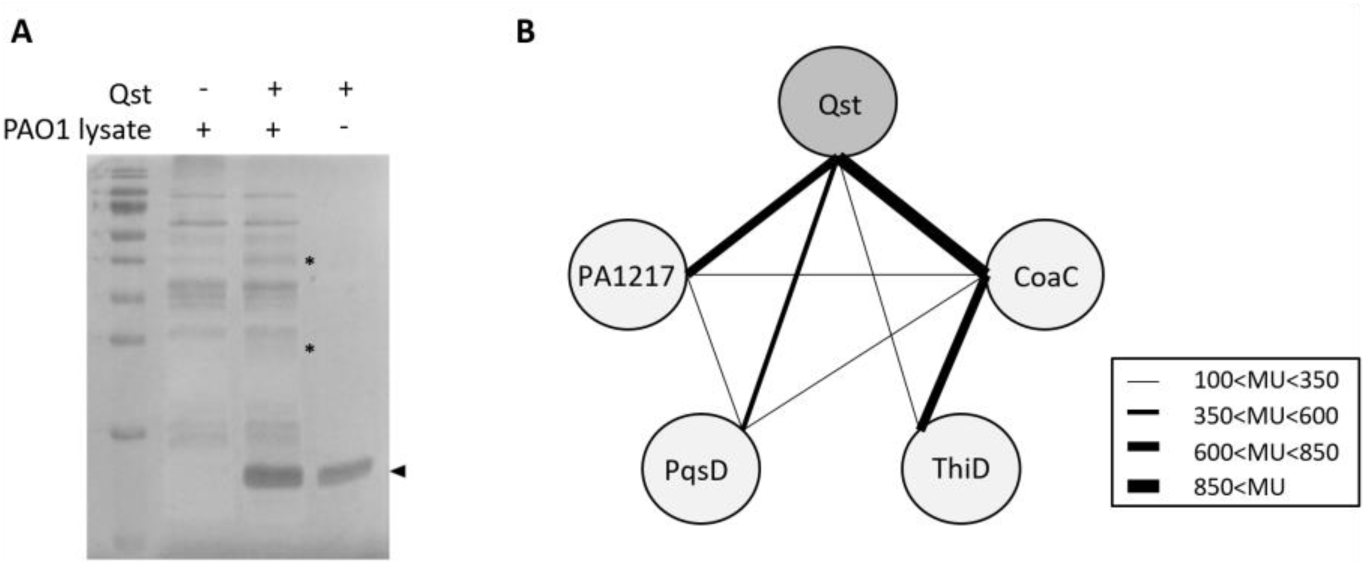
Interaction analyses of Qst and *P. aeruginosa* proteins. (A) *In vitro* pull-down of *P. aeruginosa* cell lysate, using His-tagged Qst as bait. Eluted samples were loaded on a 16% SDS-PAGE gel. The arrow indicates the band of Qst and the asterisks indicate the bands in both control and pull-down sample submitted for identification by mass spectrometry analysis. (B) Bacterial two-hybrid assay in which the T18 domain was N-terminally fused to Qst, PA1217, PqsD or ThiD and the T25 domain was N-terminally fused to PA1217, PqsD, CoaC or ThiD. Interactions were quantified by measuring the β-galactosidase activity using the Miller assay and the results (means of three biological replicates (n=3)) are visualized with black lines, MU = Miller units.

A subsequent bacterial two-hybrid experiment with Qst, PA1217, CoaC, PqsD and ThiD consolidated the identified interactions, and revealed a complex network of mutual interactions between these proteins (Figure 5B, Figure 5-figure supplement 1).

### The PQS quorum sensing system plays a key role in LUZ19 infection

A recent transcriptome analysis suggested that PQS is involved in a general host response to phage infection (Blasdel *et al.*, 2018). To further investigate the host response, we reanalyzed a metabolomics dataset of the *Pseudomonas* phage infection process (De Smet *et al.*, 2016), that reported that phage LUZ19 may target the PQS production at a post-transcriptional level. Indeed, upon LUZ19 infection, the metabolite levels of PQS were not increased, contrary to other *P. aeruginosa* infecting phages (Figure 6A, Figure 6-Source data 1). We tested several deletion mutants of the PQS biosynthesis pathway and found that a functional PqsD is important for an efficient LUZ19 infection (Figure 6B, C, Figure 6-figure supplement 1). This suggests that the interaction between Qst and PqsD has a pivotal role in host manipulation. Moreover, when supplementing the medium with HHQ, a normal LUZ19 infection in the *pqs*D deletion mutant was observed (Figure 6C), proving that LUZ19 exploits this quorum sensing system during phage infection.

**Figure 6.**
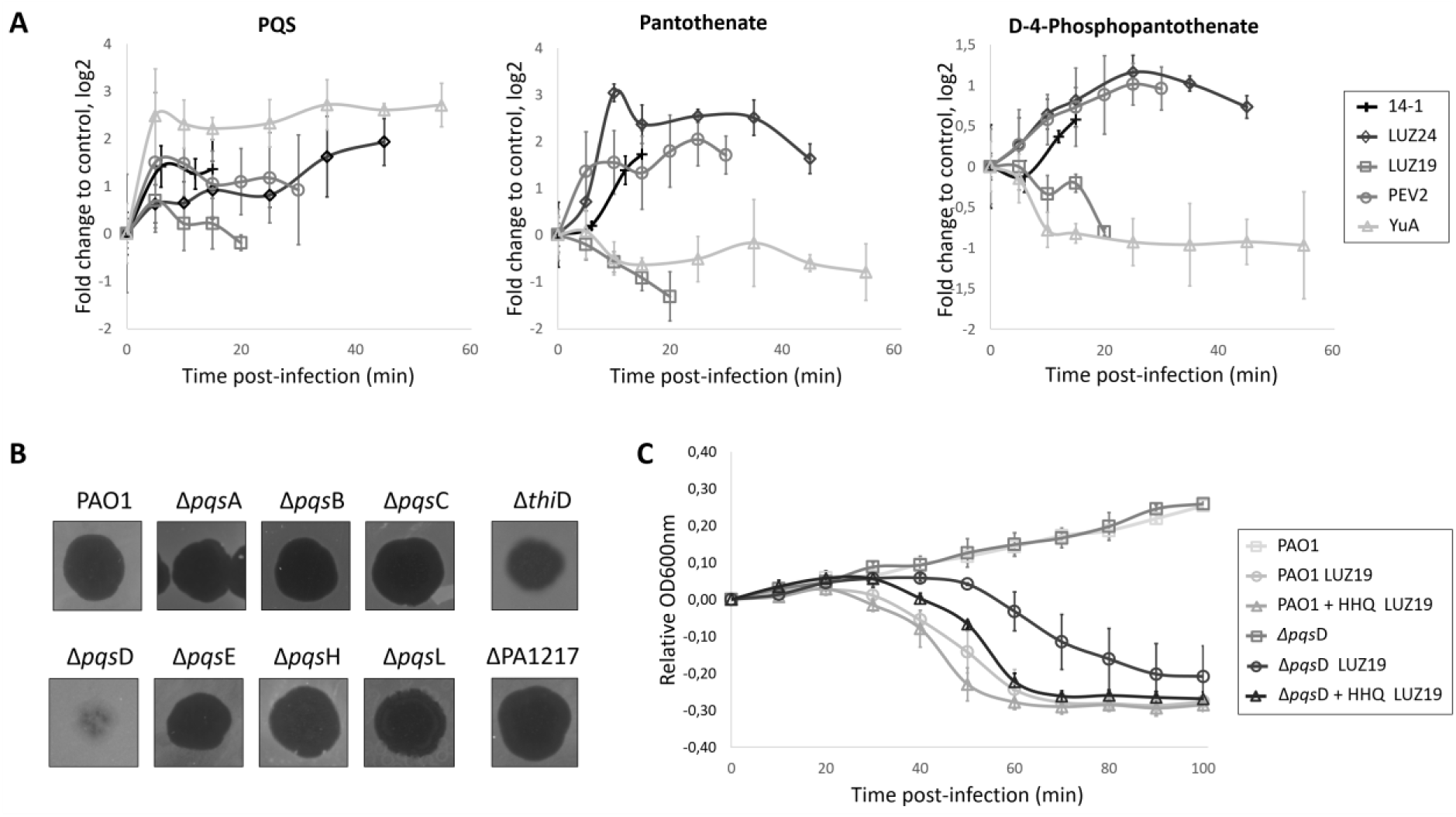
Role of Qst during LUZ19 infection. (A) Comparison of PQS, pantothenate and 4-phosphopantothenate metabolite levels in wild-type *P. aeruginosa* strains infected with different, unrelated clades of phages including 14-1, LUZ24, PEV2 and YuA (Ceyssens and Lavigne, 2010). The high-coverage metabolomics data of the *P. aeruginosa* phage infections were obtained from De Smet *et al.* (2016). Fold changes were calculated in comparison to control samples and normalized to time 0 of infection. Error bars represent fold change standard deviations (n=3). (B) Spot assay of phage LUZ19 on transposon mutants, in which approximately 10 plaque-forming units (pfu) of LUZ19 were spotted on a bacterial lawn. (C) Growth curves of *P. aeruginosa* PAO1 wild type and *pqsD* transposon mutant with or without LUZ19 infection, and supplemented with 100 µM HHQ. Exponentially growing *P. aeruginosa* cells (OD_600nm_ of 0.3) were infected with a multiplicity of infection (MOI) of 10 and followed over time. Each data point represent the mean of the OD_600nm_, normalized to time 0 of infection, of three replicates and error bars represent standard deviations (n=3).

A comparison of the metabolomes between phages-infected *P. aeruginosa* (De Smet *et al.*, 2016) and bacteria expressing Qst also supported the identified interaction with CoaC. Indeed, we observed a clear discrepancy in metabolite levels of the upstream products of CoaC (pantothenate and 4-phosphopantothenate) between LUZ19 and other *P. aeruginosa* infecting phages, all except phage YuA (Figure 6A).

## Discussion

Lytic bacteriophages have only a limited time period of a few minutes in which they hijack the host metabolism for phage production. In this process, early-expressed proteins perform specialized functions with extensive phenotypical effects on the host cell (Roucourt and Lavigne, 2009; Wagemans *et al.*, 2015, 2014). This work presents a phage-encoded quorum sensing and general host metabolic influencing protein, which was shown to be part of a previously unknown interaction network and to cause a specific inhibitory activity to the growth of *P. aeruginosa*.

### Qst targets a complex functional interaction network

By combining metabolomics and interactomics (both *in vivo* and *in vitro)*, we identified four interaction partners of Qst with high confidence. Two interaction partners are involved in the central carbon metabolism (CoaC and ThiD) and two are part of a well-studied (PqsD) and a predicted (PA1217) signaling molecule biosynthesis pathway. Since Qst, PA1217 and PqsD all share an acyltransferase activity, for which CoaC supplies the cofactor CoA and 4-phosphopantetheine (prosthetic group in acyl carrier proteins) (Kanehisa and Goto, 2000), we suggest that there is a connection between the physical interactions and functional relationships. The binary interactions between PA1217, PqsD and CoaC possibly support an efficient channeling of metabolites and might impose an additional level of regulation to these enzymes. Furthermore, the identified interaction between ThiD and CoaC is in concordance with the known connection between the two vitamin-associated pathways they are part of, namely of thiamine and CoA respectively (Enos-Berlage and Downs, 1997; Frodyma *et al.*, 2000). The fact that ThiD does not show an interaction neither with PA1217 nor with PqsD further supports our results, since no relationships between these enzymes are reported. Interestingly, the interactions CoaC-PqsD and CoaC-ThiD were predicted using a machine learning-based integrative approach (Zhang *et al.*, 2012).

Qst targets multiple enzymes and thereby affects multiple metabolic pathways, including two central metabolic pathways and quorum sensing. This broad impact may explain the growth-inhibitory effect of Qst on *P. aeruginosa* cells and highlights the extent of evolutionary molecular adaptation by LUZ19 to its host. It is unclear how Qst influences these enzymes. However, the interaction between Qst and the catalytic part of PA1217 indicates a potential influence of Qst on the enzymatic activity or specificity of its targets. Also the direct effect of Qst on the substrate and product levels of CoaC and ThiD respectively, suggests a phage-mediated overactivation of these enzymes. Since all interaction partners are linked with each other forming a functional network, it can be hypothesized that Qst has a regulatory function, possibly through post-translational modification or direct protein-protein interactions.

### Metabolic changes mark Qst as an influencer of the PQS-mediated host response to phage infection

One of the key observations is the reduction of metabolite levels in *P. aeruginosa* of the downstream products of PqsD (DHQ, HHQ and PQS) when recombinant Qst is present in the cell. To our knowledge, this is the first report of a phage protein influencing PQS. Targeting PQS can have a major impact on the host cell’s physiological state, as PQS is one of the main quorum sensing systems in *P. aeruginosa*, implicating that it impacts pathogenicity but also general metabolic functions like cofactor biosynthesis and fatty acid metabolism (Häussler and Becker, 2008; Schuster and Greenberg, 2006). Besides quorum sensing signaling, PQS has also been implicated in iron acquisition, cytotoxicity, outer-membrane vesicle biogenesis, and modulation of the host immune response (Lin *et al.*, 2018), making it a key regulatory hub to stress conditions, for example during phage infection.

Previous transcriptomics data from our lab indicates that PQS is responsible for a general host response to phage infection, by significantly upregulating the *pqsABCDE* operon and the co-regulated *phnAP* operon during infection of several distinct lytic *P. aeruginosa* infecting phages (Blasdel *et al.*, 2018). Despite a general PQS transcription response to phage infection, an increase of metabolite levels has not been observed for phage LUZ19 contrary to other *P. aeruginosa* infecting phages (De Smet *et al.*, 2016). This was the first indication that LUZ19 may encode a protein which prevents PQS production on the post-transcriptional level or exploits this system to support its infection, and our results reveal that Qst is the responsible protein. Since a functional PqsD enzyme is important for efficient LUZ19 infection, it can be hypothesized that LUZ19 not specifically tackles host response but rather uses the PQS biosynthesis system to create a favorable environment for phage production, for example by changing the PqsD activity or its products through acylation by a still unknown mechanism. Indeed, external supplementation of the quorum sensing molecule to *P. aeruginosa* cells lacking PqsD restored normal LUZ19 infection.

PA1217 is also part of a predicted signaling molecule biosynthesis pathway, namely the NRPS cluster PA1221 (Gulick, 2017). It can therefore also be hypothesized that there is a link between these two cell-to-cell signaling systems, which produce stress-related metabolites, suggesting a host response to phage infection. Indeed, PA1217 and related proteins belonging to the same cluster (PA1218 and PA1221) are also influenced during phage infection of phages PaP1 (*Myoviridae*) and PaP3 (*Podoviridae*), respectively (Zhao *et al.*, 2017, 2016). Future work will aim at identifying and characterizing this secondary metabolite to unravel its function in *P. aeruginosa* and its role during phage infection, as a potentially extra quorum sensing mechanism.

### Qst can be a valuable study tool in the battle against antibiotic resistant *P. aeruginosa*

The use of quorum sensing by bacteria to resist phage infection has recently been suggested (Hoyland-Kroghsbo *et al.*, 2017, 2013; Patterson *et al.*, 2016; Qin *et al.*, 2017). However, the opposite, in which phages target quorum sensing to counter the bacterial defense mechanism or to create a favorable environment for phage infection has, to our knowledge, never been observed. The fact that Qst targets PqsD is even more intriguing, as this enzyme catalyzes the last and key step in HHQ biosynthesis, which make it a widely studied anti-virulence and anti-biofilm target to combat *P. aeruginosa* infections (Storz *et al.*, 2012; Weidel *et al.*, 2013; Zhou and Ma, 2017). Furthermore, the Qst interaction partners CoaC and ThiD are studied for their antibacterial drug target potential, as they are different in eukaryotic cells and inhibition will lead to the depletion of vitamins, thereby damaging the growth of cells (Du *et al.*, 2011; O’Toole and Cygler, 2003).

## Material and methods

### Phages and bacterial strains

*Pseudomonas aeruginosa* PAO1 and derivatives were used in this study (Stover *et al.*, 2000). Strain PAO1-R1, a lasR^−^lasI^−^ mutant containing the pTS400 plasmid (*lasB’-lacZ* translational fusion), was used for the PQS bioassay (Gambello and Iglewski, 1991). Transposon deletion mutants were ordered from the *P. aeruginosa* mutant library (Jacobs *et al.*, 2003): PW2799, PW2800, PW2803, PW2805, PW2806, PW5343, PW8105, PW7728, PW3198. A *P. aeruginosa* PAO1 strain with single-copy integration of the phage gene under a lac promotor into the bacterial genome was made using the Gateway cloning system (Invitrogen, Carlsbad, CA), as described previously (Hendrix *et al.*, 2019). Three *Escherichia coli* strains were used: *E. coli* TOP10 (Life Technologies, Carlsbad, CA) for cloning procedures, *E. coli* BTH101 (Euromedex, Souffelweyersheim, F) for Bacterial two-hybrid assays and *E. coli* BL21(DE3) (Life Technologies) for recombinant protein expression.

### P. aeruginosa PAO1 genome fragment library construction

A *P. aeruginosa* PAO1 fragment library was constructed by ligating genomic DNA fragments into the pHERD20T vector (Qiu *et al.*, 2008). For this, restriction enzymes AluI and RsaI (Thermo Scientific, Waltham, MA) were applied to fragment 37.5 µg purified genomic DNA, and restriction enzyme SmaI (Thermo Scientific) was used to digest 3 µg pHERD20T vector according the manufacturer’s protocol. The fragments with lengths between 1.5 kbp and 4 kbp were isolated using gel extraction and concentrated by ethanol precipitation. The digested vector, on the other hand, was treated with alkaline phosphatase FastAP (Thermo Scientific). The ligation reaction mixture, with a total volume of 20 µl, contained a three-fold excess of DNA fragments compared to the pHERD20T vector (50 ng), 1 µl T4 DNA ligase, 2 µl T4 buffer and 2 µl 50% PEG4000 (Merck, Kenilworth, NJ). After incubation, the entire library was transformed to chemically competent *E. coli* TOP10 cells and the number of transformants was determined.

### Complementation assays

The *P. aeruginosa* PAO1 genome fragment library, containing random fragments ranging from 1.5 to 4 kbp inserted in a pHERD20T vector under control of a p*BAD* promotor, was electroporated (Choi *et al.*, 2006) to the mutant *P. aeruginosa* PAO1 strain encoding *qst*. Cells were grown overnight at 37°C on LB agar supplemented with gentamicin (30 µg/ml), carbenicillin (200 µg/ml), 1 mM isopropyl-β-D-1-thiogalactopyranoside (IPTG) and 0.2% L-arabinose (L-ara). Ninety-six colonies were selected and checked for false positive hits by repeating the growth step on selective media. The plasmids of the positive hits were isolated using the GeneJET Plasmid Miniprep Kit (Thermo Scientific) and transformed afresh to the *P. aeruginosa qst* strain to verify if the phenotype is indeed due to the presence of the fragment. After confirmation, a single colony was picked to perform a spot test. From each overnight culture, a hundredfold dilution series was spotted in triplicate on LB agar with or without IPTG/L-ara and overnight incubated at 37°C. To determine the location of the fragments in the *P. aeruginosa* genome, the sequencing results were analysed with Basic Local Alignment Search Tool (BLAST; NCBI (Altschul *et al.*, 1990)) and the *Pseudomonas* Genome Database (Winsor *et al.*, 2016). Individual genes, which were located in the maximal overlap region of the identified fragments, were cloned in the pME6032 vector (Heeb *et al.*, 2002) for electroporation to the *P. aeruginosa qst* strain. Again, a dilution series was spotted on LB with and without 1 mM IPTG and incubated overnight at 37°C. As a negative control, the empty pME6032 vector was used.

### Time-lapse microscopy

Microscopic analysis was performed according to a previously described procedure (Wagemans *et al.*, 2014). Briefly, an overnight culture of *P. aeruginosa* cells, containing a single-copy *qst* expression construct and the multicopy expression vector pME6032 with or without bacterial gene, was diluted thousand times and spotted on LB agar supplemented with gentamicin (30 µg/ml), tetracycline (60 µg/ml) and 1 mM IPTG. Time-lapse microscopy was performed for 5 h at 37°C with a temperature controlled Nikon Eclipse Ti time-lapse microscope using the NIS-Elements AR 3.2 software (Cenens *et al.*, 2013).

### Protein expression and purification

Qst was fused to a His-tag using the pEXP5-CT/TOPO vector (Invitrogen) following the TA-cloning protocol provided by the manufacturer. The bacterial protein PA1217 was fused to a GST-tag by cloning it into a pGEX-6P-1 vector (GE Healthcare, Chicago, IL) using the BamHI and EcoRI restriction sites (Thermo Scientific). Recombinant expression of both proteins was performed in exponentially growing *E. coli* BL21(DE3) cells after induction with 1 mM IPTG at 30°C overnight. The proteins were purified using a 1 ml HisTrap HP column (GE Healthcare) or 5 ml GSTrap HP column (GE Healthcare), depending on the fused tag, on an Äkta Fast Protein Liquid Chromatography (FPLC, GE Healthcare) system, followed by dialysis to storage buffer (50 mM Tris-HCl, pH 7.4, 150 mM NaCl) supplemented with 5% (PA1217) or 15% (Qst) glycerol and stored at −20°C.

### In vitro pull-down

A total of 250 ng of Qst was incubated with 1 ml Ni-NTA Superflow beads (Qiagen, Hilden, DE) in an end-over-end shaker for 1 h at 4°C. The supernatant was removed by centrifugation at 300 *g* for 5 min and the mixture was suspended in 10 ml pull-down buffer (20 mM Tris pH 7.5, 200 mM NaCl, 20 mM imidazole). The Ni-NTA matrix was loaded on a prepared 1 ml Polypropylene column (Qiagen) and a filtered *P. aeruginosa* cell lysate was added. For the bacterial lysate, cells were grown in 1 L LB until an OD_600nm_ of 0.6, pelleted and resuspended in 40 ml protein A buffer (10 mM Tris pH 8.0, 150 mM NaCl, 0.1 % (v/v) NP-40) supplemented with 40 mg HEWL and 24 mg Pefabloc®SC. Cell lysis was done by three freeze-thaw cycles and sonication (40% amplitude, 8 times 30 sec with 30 sec in between). The column was washed three times with pull-down buffer, after which 500 µl pull-down buffer supplemented with 500 mM imidazole was added. After briefly vortexing the column and 5 min incubation at room temperature, the proteins were eluted by centrifugation at 1,000 *g* for 5 min and visualized using SDS-PAGE. The protein bands that were present in the pull-down sample but absent in the control sample (without loaded Qst) were analyzed using mass spectrometry analysis as described previously (Van den Bossche *et al.*, 2016).

### Native mobility shift assay

Interaction between the phage and bacterial protein was evaluated *in vitro* using a native mobility shift assay (Van den Bossche *et al.*, 2014). Mixtures of 200 µM *P. aeruginosa* PA1217 and an increasing amount of Qst were incubated in reaction buffer (20 mM Tris, pH 8) for 10 min at room temperature (T_R_). After the addition of loading dye (0.2% (w/v) bromophenol blue, 300 mM DTT and 50% (v/v) glycerol), the samples were loaded on a 10% polyacrylamide native gel (10% (v/v) 37.5:1 acrylamide/bisacrylamide, 10% (v/v) glycerol, 200 mM Tris, pH 8, 0.01% (v/v) APS, 0.001% (v/v) TEMED) and run in running buffer (25 mM Tris, 250 mM glycine) for 80 min at 150 V. The gel was Coomassie-stained, the shifted bands were excised from the gel and the gel pieces were loaded on a 12% SDS-PAGE gel to confirm the presence of both proteins. Furthermore, the proteins were identified using mass spectrometry analysis as described previously (Van den Bossche *et al.*, 2016).

### Enzyme-linked immunosorbent assay (ELISA)

ELISA was performed using anti-GST coated strips (Pierce™ Anti-GST Coated Clear Strip Plates, Thermo Scientific). A fixed amount of 10 pmol GST-tagged PA1217 was diluted in 200 µl PBS (137 mM NaCl, 2.7 mM KCl, 8.2 mM Na_2_HPO_4_, 1.8 mM KH_2_PO_4_, pH 7.5) supplemented with 2% (w/v) Bovine serum albumin (BSA), added to the wells and incubated for 1 h at T_R_ while gently shaking. After washing the wells three times with PBS-Tween 0.1% followed by three times PBS, increasing amounts of His-tagged Qst were diluted in 200 µl PBS + 2% BSA, added to the wells and incubated for 1 h at T_R_. The washing step was repeated and 200 µl of a 1:5,000 dilution of the monoclonal Anti-His antibody (Sigma-Aldrich, St. Louis, MO) in PBS + 2% BSA was added to each well. After incubating 1 h at T_R_, the wells were washed and 200 µl of a 1:5,000 dilution of secondary Anti-Mouse IgG antibody conjugated to HRP (Promega Corporation, Madison, WI) in PBS + 2% BSA was added to each well. A final washing step after 1 h incubation at T_R_ was followed by addition of 100 µl 1-Step Slow TMB-ELISA substrate (Thermo Scientific) to each well and absorbance was measured after 10 min incubation at T_R_ using the Bio-Rad Model 680 Microplate Reader (655 nm; Bio-Rad, Hercules, CA). As negative controls, wells without bait protein (GST-tagged PA1217) and pure GST tag were used. Three replicates of each combination were tested

### Bacterial two-hybrid

Bacterial two-hybrid assays were conducted using the BACTH System kit (bacterial adenylate cyclase two-hybrid system kit, Euromedex). The *P. aeruginosa* genes *PA1217, pqsD* and *thiD* were cloned into four vectors (pUT18, pUT18C, pKT25 and pN25), while the fragments of the former (sequences encoding amino acid 1-259, 1-342, 97-317, 156-342 and 260-455) were only cloned into pKT25. The *P. aeruginosa* genes *coaC, pqsB* and *pqsC* were cloned into vectors pKT25 and pN25, and the phage gene *qst* was cloned into the vectors pUT18, pUT18C and pN25. Each combination of phage and bacterial gene/fragment was co-transformed to *E. coli* BTH101 and dilutions of overnight cultures were spotted on synthetic minimal M63 medium (15 mM (NH_4_)_2_SO_4_, 100 mM KH_2_PO_4_, 1.7 µM FeSO_4_, 1 mM MgSO_4_, 0.05% (w/v) vitamin B1, 20% (w/v) maltose) supplemented with 0.5 mM IPTG and 40 µg/ml 5-bromo-4-chloro-3-indolyl-β-D-galactopyranoside (X-gal). To quantify the β-galactosidase activity, Miller assays were performed (Zhang and Bremer, 1995). As negative controls, the constructs were co-transformed with their empty counterparts. Each combination was tested in triplicate.

### CoA acetyltransferase activity assay

To measure acetyl-CoA dependent acetyltransferase activity of the bacterial and phage protein, an end-point assay was performed as previously described (Roucourt *et al.*, 2009). Briefly, the formation of CoA was determined using 5,5’-dithiobis-(2-nitrobenzoic acid) (DTNB), which reacts with free thiol groups of CoA. Mixtures of 1 µg PA1217 and/or 5 µg Qst, 2 mM 3-methyl-2-oxobutanoic acid (Fisher Scientific, Hampton, NH) and 1 mM acetyl-CoA (Sigma-Aldrich) were incubated in a reaction buffer (70 mM Tris, pH7.5, 3.5 mM MgCl_2_, 3.5 mM KCl) for 30 min at 37°C. After the addition of 2 mM DTNB (Sigma-Aldrich) to each sample, the absorption was measured at 415 nm in a microplate reader (Microplate reader model 680, Bio-Rad). As a positive and negative control, samples with LeuA and without protein and pure GST tag were tested, respectively. The experiment was performed in triplicate.

### ^1^H-NMR analysis

The predicted α-isopropylmalate synthase activity of PA1217 was studied using ^1^H-NMR (de Carvalho and Blanchard, 2006). Mixtures of 100 µg PA1217, 12 mM MgCl_2_, 1.1 mM 3-methyl-2-oxobutanoic acid and 1 mM acetyl-CoA were incubated in 50 mM potassium phosphate buffer (pH 7.5) for 2 h at 37 °C. The proteins were removed using Amicon Ultra 3K 0.5 ml centrifugal filters (Millipore, Burlington, MA) according to the manufacturer’s protocol. Lastly, 300 µl of D_2_O was added to 300 µl of the prepared mixture. ^*^1^*^*H-NMR* spectra were recorded on a Bruker Ascend 600 MHz and 400 MHz spectrometer equipped with a BBO 5 mm atma probe and a sample case. To suppress the broad signal of water, an adapted zgpr pulse program (p1 9.75 µs; plw1 15W; plw9 5.7-05W; o1P) was applied on the resonance signal of water, which was determined and selected automatically. The experiment was repeated with mixtures containing Qst, a combination of Qst and PA1217, and the positive control LeuA.

### Metabolomics using FIA-qTOF-MS

Metabolites were extracted from *P. aeruginosa* cells and analyzed following the procedure described previously (De Smet *et al.*, 2016). Cells were grown until an OD_600nm_ = 0.1, 1 mM IPTG was added to each culture and sampling was done every 15 min. Four strains were tested: PAO1 wild type, PAO1 encoding *qst*, PAO1 containing the multicopy expression vector pME6032 with *PA1217* gene, and PAO1 combining the previous two strains. The samples were analysed using negative mode flow injection-time-of-flight mass spectrometry (FIA-qTOF). Ions were annotated with KEGG *P. aeruginosa* metabolite lists (Kanehisa and Goto, 2000), allowing a mass tolerance of 1 mDa and an intensity cutoff of 5000 counts. Raw intensities were quantile normalized (each measurement is an average of two technical replicates) and fold changes were calculated for each time point between each condition and control for means of three biological replicates and reported in log2. *P*-values were calculated with unpaired t-test and corrected for multiple hypothesis testing with Benjamini-Hochberg procedure. Page’s trend test p-values were calculated using the Matlab Page package: http://www.mathworks.com/matlabcentral/fileexchange/14419-perform-page-test?focused=6141676&tab=function.

### PQS bioassay

The effect of the phage and bacterial protein on the production pattern of PQS autoinducer by *P. aeruginosa* was tested using the *P. aeruginosa* PAO1-R1 (pTS400) reporter strain (Van Houdt *et al.*, 2004). First, cell-free culture supernatants from tested *P. aeruginosa* strains were prepared as described by Van Houdt *et al.* (2004) with some modifications. Briefly, overnight cultures of *P. aeruginosa* cells (over)expressing the *qst* and/or the PA1217 gene were diluted (1:100) in fresh LB medium and grown for 21 h at 37°C. The cells were then removed by centrifugation (24,000 g) for 10 min and cleared supernatants was filtered using 0.22 µm Millex filter units (Millipore). Next, 2 ml of the cell-free culture supernatant was mixed with 2 ml of freshly diluted (1:100) *P. aeruginosa* PAO1-R1 (pTS400) culture in PTSB medium (5% (w/v) peptone, 0.25% (w/v) trypticase soy broth), and incubated for 18 h at 37°C. The production of reporter enzyme β-galactosidase was measured using previously mentioned Miller assay (Zhang and Bremer, 1995). The experiment was performed in triplicate.

## Acknowledgements

We would like to thank prof. Dirk De Vos (Centre for Surface Chemistry and Catalysis, KU Leuven, Belgium) for use of the NMR facilities, and Erik Royackers (Hasselt Univeristy, Belgium) for the technical support in mass spectrometry analysis. This research was supported by the KU Leuven project GOA ‘Phage Biosystems’.

## Figure supplements

**Figure 2-figure supplement 1.**
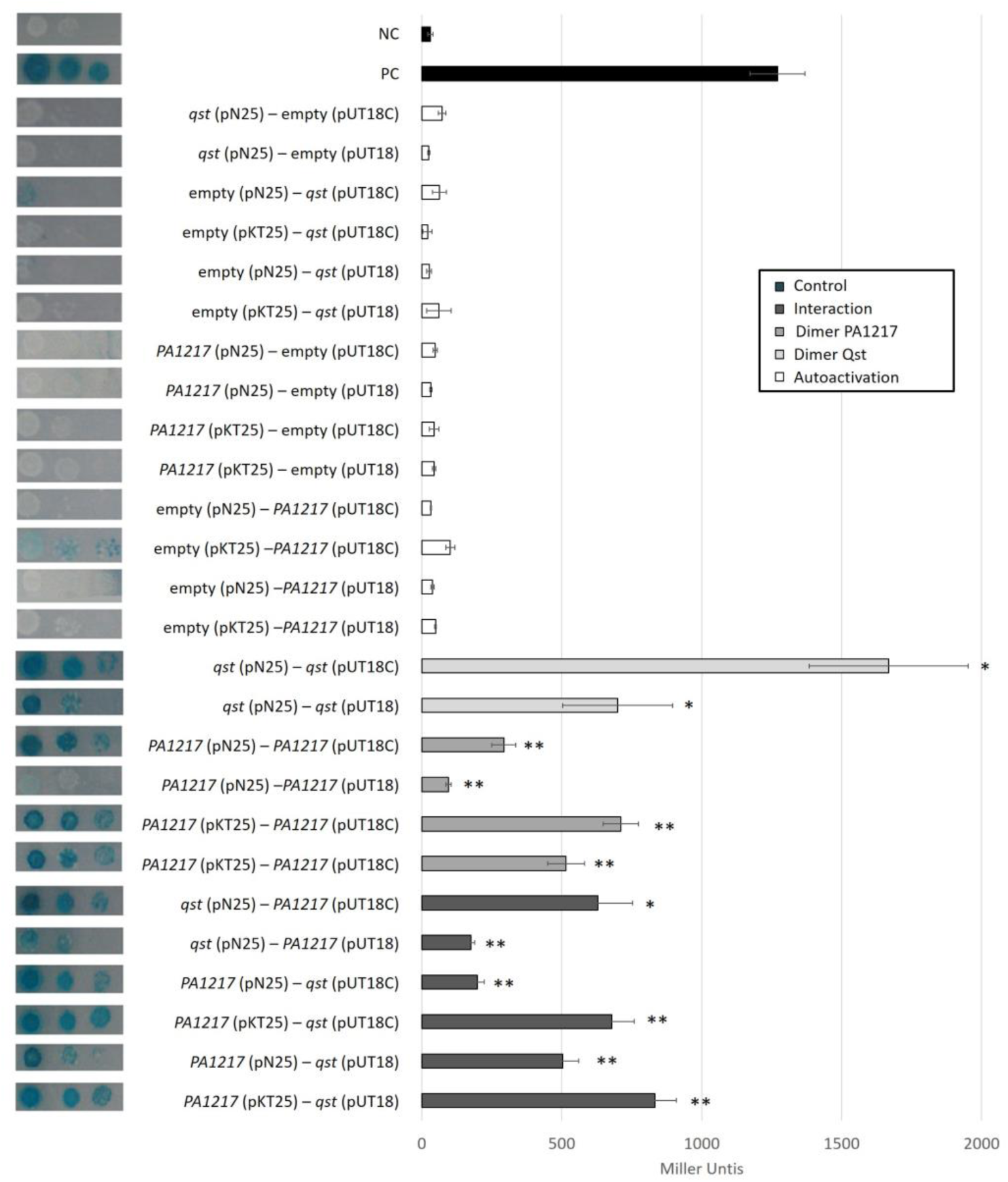
Bacterial two-hybrid results of Qst and PA1217. Interactions were visualized by a drop test on minimal medium, and quantified by measuring the β-galactosidase activity, which are indicated in Miller Units. Non-fused T25 and T18 domains were used as a negative control, and the leucine zipper of GCN4 was used as a positive control. Error bars represent standard deviation and *P*-values (compared to both controls) were calculated using the Student’s t-test (n=3), *p<0.5, **p<0.01.

**Figure 3-figure supplement 1.**
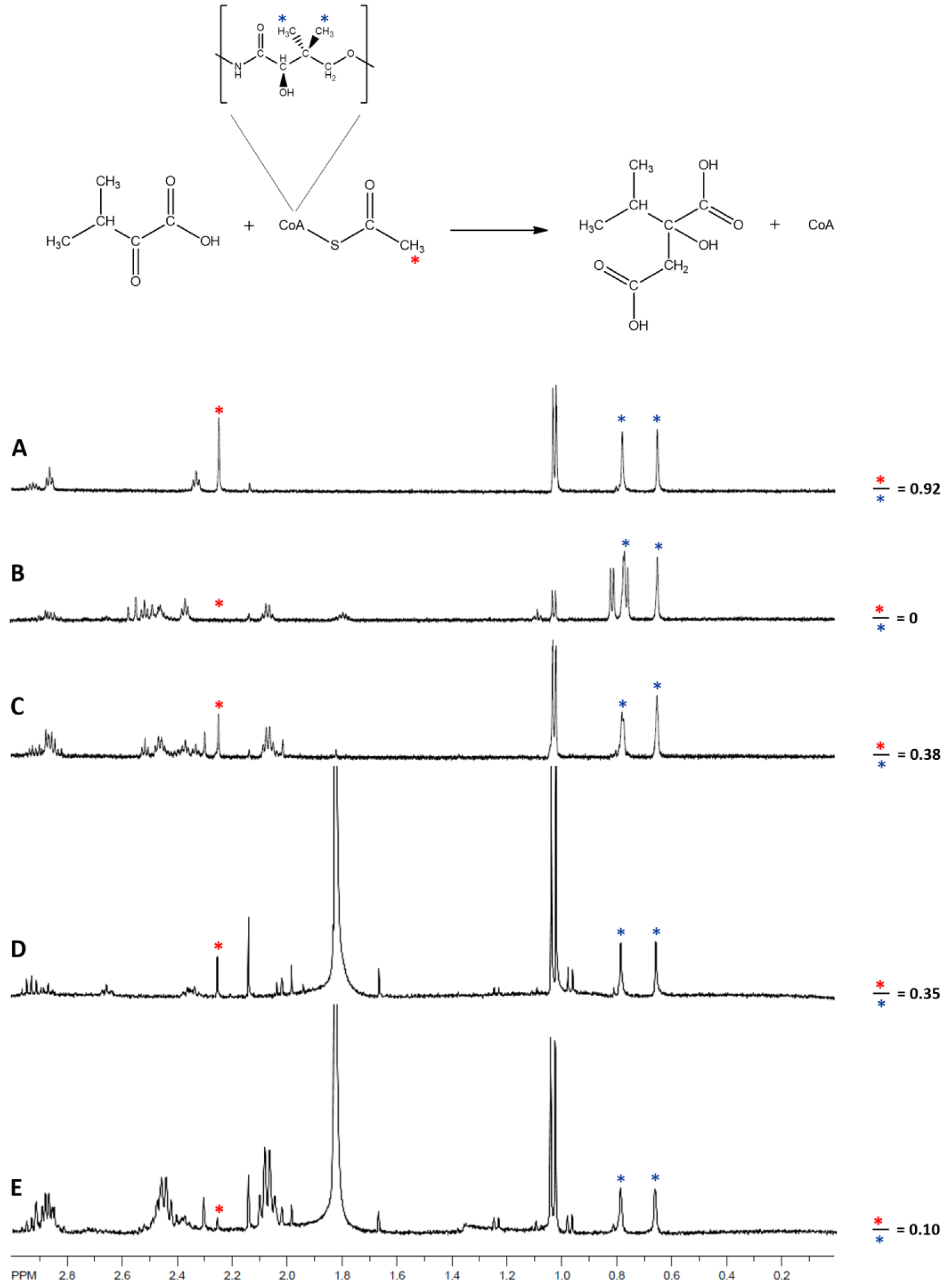
^***^1^***^***H*-NMR analysis of acetyltransferase and 2-isopropylmalate synthase activity**. Region of spectral interest of the reaction of the 2-isopropylmalate synthase LeuA (B), the hypothetical protein PA1217 (C), Qst (D) and both PA1217 and Qst (E) with 3-methyl-2-oxobutanoate and acetyl-CoA. The top spectrum (A) shows the reaction mixture without enzyme. The ratio of peak 4 compared to peak 5 shows the percentage of intact acetyl-CoA molecules in the reaction mixture. The lower the amount, the more acetyl-CoA molecules were used by the protein.

**Figure 4-figure supplement 1.**
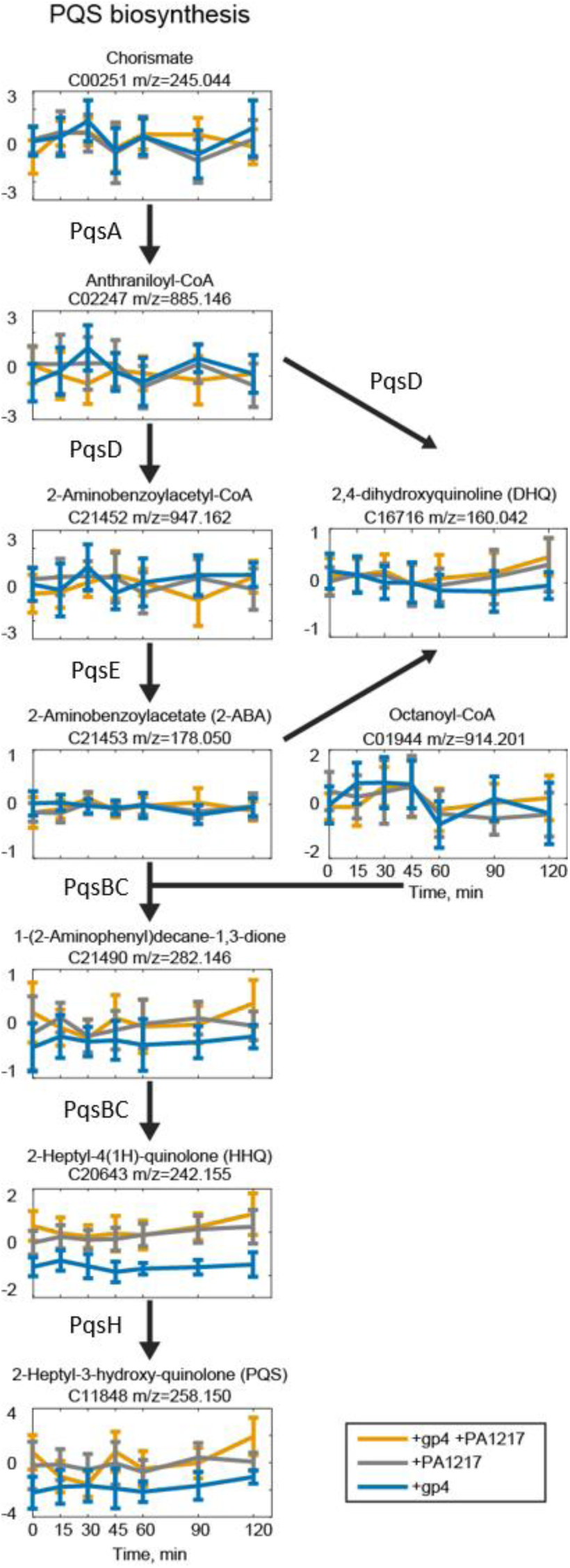
Metabolomic results of the *P. aeruginosa* quinolone signal (PQS) pathway after Qst and PA1217 expression. Graphs show the fold changes of metabolites in the PQS pathway compared to IPTG-induced wild-type *P. aeruginosa* PAO1 at different stages of expression (0, 15, 30, 45, 60, 90 and 120 min). The x axis shows the time points during infection (in minutes), and the y axis shows the fold changes compared to the control. Annotated metabolite name and ion mass are shown above the graph. Error bars represent fold change standard deviations (n=3).

**Figure 5-figure supplement 1.**
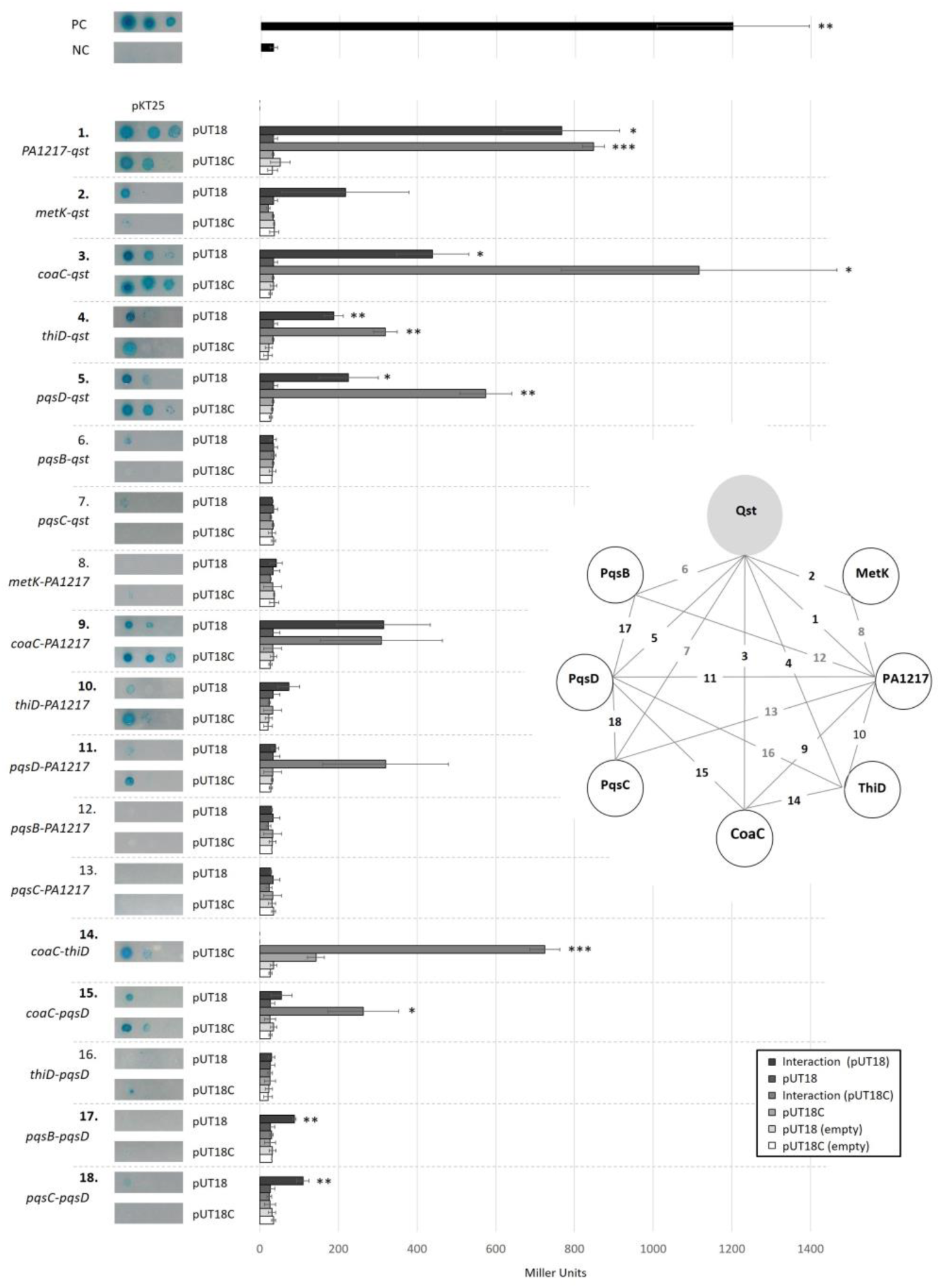
Bacterial two-hybrid results of Qst with PA1217, MetK, CoaC, ThiD, PqsB, PqsC and PqsD. The diagram shows the tested combinations, in which the numbers (black: interaction; grey: no interaction) correspond with the numbers in the graph. The T25 domain is N-terminally fused to the first listed protein, the T18 domain is C-(pUT18) or N-terminally (pUT18C) fused to the second protein. Interactions were visualized by a drop test on minimal medium, and quantified by measuring the β-galactosidase activity, which are indicated in Miller Units. Non-fused T25 and T18 domains were used as a negative control, and the leucine zipper of GCN4 was used as a positive control. Error bars represent standard deviation and *P*-values (compared to both controls) were calculated using the Student’s t-test (n=3), *p<0.5, **p<0.01, *** p<0.001.

**Figure 6-figure supplement 1.**
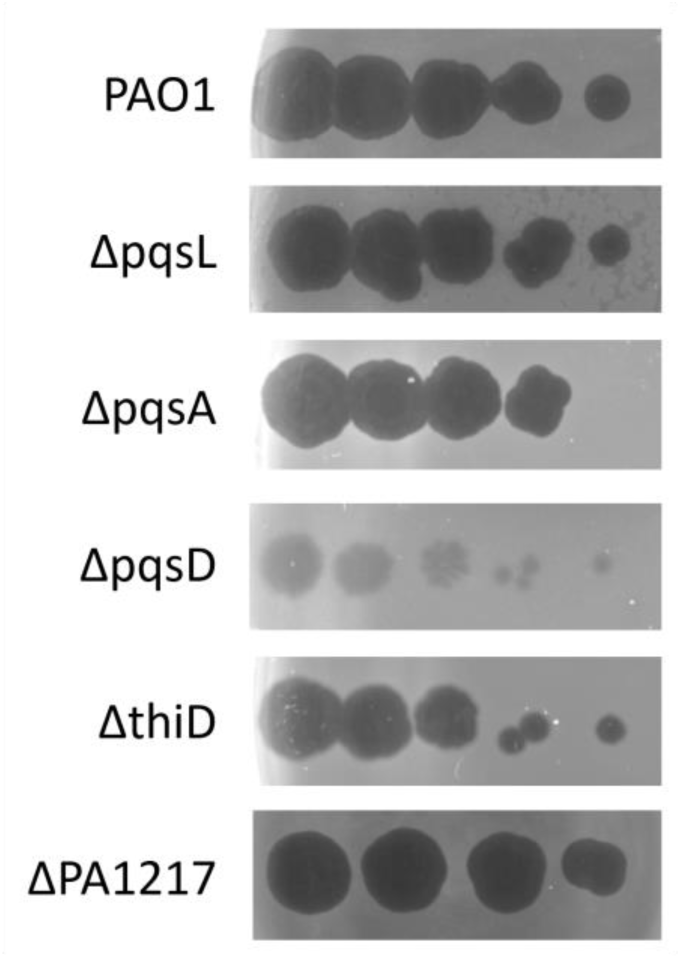
Serial dilutions of phage LUZ19 on wild-type *P. aeruginosa* PAO1 (top) and transposon mutants. One microliter of, from left to right 10^7^, 10^6^, 10^5^, 10^4^ and 10^3^ pfu/ml, dilution was spotted on lawns of (mutant) *P. aeruginosa* strains.

## Source Data

**Figure 2-Source Data 1. Interaction analyses between Qst and PA1217**

**Figure 2-figure supplement 1-Source data 1. Bacterial two-hybrid results of Qst and PA1217**

**Figure 3-Source Data 1. CoA acetyltransferase activity assay**

**Figure 4-Source Data 1. Metabolites influenced in *P. aeruginosa* expressing Qst.**

Levels of metabolites in wild-type *P. aeruginosa* PAO1 strain (control), *P. aeruginosa* PAO1 strains expressing Qst (gp4), overexpressing PA1217 (PA1217), and in a strain both expressing Qst and PA1217 (double) are given. For each condition and control, the normalized ion intensities, ion fold changes, fold change standard deviations, p-values calculated with unpaired t-test for each time point, and Page’s trend test p-values for the time-course data are showed.

**Figure 4-Source Data 2. PQS bioassay**

**Figure 5-Source Data 1. *In vitro* pull down of *P. aeruginosa* cell lysate, using His-tagged Qst as a bait.**

MS results of the *in vitro* pull-down with His-tagged Qst as bait and *P. aeruginosa* cell lysate as prey. The numbers in the summarized table indicate the ‘Total spectral Count’ identified for a specific protein. Analyses were done on a LTQ-Orbitrap Velos Pro (Thermo Scientific).

**Figure 5-figure supplement 1-Source Data 1. Bacterial two-hybrid results of Qst, PA1217, MetK, CoaC, ThiD, PqsB, PqsC and PqsD**

**Figure 6-Source Data 1. Metabolomics of phage-infected *P. aeruginosa*.**

The high-coverage metabolomics data of the *P. aeruginosa* phage infections are obtained from De Smet *et al.* (2016). For this, fold changes were calculated in comparison to control samples and normalized to time 0 of infection.

